# Crosslinked CXCR4 Signals Decreased Motility and Increased Adhesion of T Cells

**DOI:** 10.1101/2025.06.04.652236

**Authors:** Jordan Pearson, Jung-in Yang, Sean Krivitsky, Z Kelley, Douglas T. Fearon

**Affiliations:** Cancer Center, Cold Spring Harbor Laboratory, Cold Spring Harbor, NY 11724; Graduate Program in Genetics, Stony Brook University, Stony Brook, NY 11794; Medical Scientist Training Program, Stony Brook University Renaissance School of Medicine, Stony Brook University, Stony Brook, NY 11794; Department of Biochemistry and Cell Biology, Stony Brook University, Stony Brook, NY 11794; Meyer Cancer Center, Weill Cornell Medicine, New York, NY 10065

**Keywords:** Pancreatic ductal adenocarcinoma, T cell migration, CXCL12, PTK2B, TNFα-TNFRSF1B signaling

## Abstract

Pancreatic ductal adenocarcinoma (PDA) is a highly aggressive cancer known for its ability to evade immune surveillance, primarily through mechanisms that prevent T cell infiltration. While the coating of cancer cells with the chemokine, CXCL12, is required for the exclusion of T cells, the precise molecular mechanisms behind the failure of T cell migration into the tumor remain unclear. In this study, we identify a potential mechanism by demonstrating that crosslinking CXCR4 is associated with decreased motility of T cells secondary to their adhesion to fibronectin. Using human lymphoblastoid T cells and primary human T cells, we show that polymeric CXCL12-induced crosslinked CXCR4 triggers the non-G α*i*-dependent pathway of tyrosine phosphorylation of FAK-related proline-rich tyrosine kinase 2 (PTK2B) in T cells. This response is necessary for the decreased motility and increased adhesion of T cells. We also find that the downstream cellular reactions of this pathway is secondary to CXCR4-crosslinking and require the integrin subunit, α4, and TNFα stimulation of TNFRSF1B. These findings provide insights into the mechanisms mediating the exclusion of T cells from nests of PDA cells and further support therapeutic strategies aimed at blocking the interaction of CXCR4 on T cells with the CXCL12-coating of PDA cells.

## Background

The resistance of pancreatic adenocarcinoma to immunotherapy has been ascribed to various factors. The expression of inhibitory receptors on T cells, such as PD-1 and LAG-3, can dampen T cell attack of cancer cells, but administration of blocking antibodies to these inhibitory receptors has not benefited patients or mice with PDA (1). The cancer is considered to be relatively non-immunogenic, but ongoing adaptive immune responses that lead to killing of cancer cells have been noted when a CXCR4 inhibitor, AMD3100, is continuously given to patients and mice with PDA (2, 3). Finally, T cells that mediate anti-tumor immunity have been proposed to be non-functional, or “exhausted”, but this explanation for resistance of carcinomas to immune attack would seem to be excluded by report that T cells from the draining lymph nodes are required for checkpoint inhibitor immunotherapy (4).

The exclusion of T cells from nests of carcinoma cells and restriction to the stromal regions of tumors has been recognized for over a decade (5–7). Two explanations are possible: restricted movement caused by cancer cells or trapping by stromal cells. Although support for the latter hypothesis was seemingly provided by the finding that conditionally depleting cancer-associated fibroblasts (8) induced immunological control of mouse PDA (2), this was demonstrated to be caused by the depleting the major biosynthetic source of CXCL12 that coats PDA cells and mediates the exclusion of T cells (9). Keratin-19 (KRT19) secreted by PDA cancer cells (10) remains associated with the plasma membrane and binds CXCL12, forming a polymeric CXCL12-KRT19 coating. The polymerized CXCL12-KRT19 coat is stabilized by transglutaminase-2 (TGM2), which covalently links the complex (10). PDA tumors formed with cancer cells lacking either KRT19 or TGM2 are not coated with CXCL12 and are infiltrated by T cells. Thus, the polymeric CXCL12 from this complex, in contrast to the monomeric form of CXCL12, plays a pivotal role in T cell exclusion by engaging with T cells in a distinct manner that effectively impedes their infiltration into tumor nests.

CXCR4 is the only T cell receptor that recognizes CXCL12, as the T cell does not express ACKR3 (11). The recognized signaling function of CXCR4, which is to mediate the chemotactic responses to monomeric CXCL12, cannot explain the phenomenon of CXCL12/CXCR4-dependent T cell exclusion. However, a broader regulatory function with respect to leukocyte movement has been demonstrated by the requirement for the interaction of CXCR4 with CXCL12 in the process of bone marrow retention of hematopoietic stem cells and other immature leukocytes (12). Furthermore, the inhibition by monomeric CXCL12 of the chemotaxis of T cells mediated by other chemokine receptors is additional evidence for a suppressive role for CXCR4 on T cell motility (3). Thus, while it is not unprecedented that CXCR4 functions include the restriction of immune cell motility, we do not have an understanding of the signaling pathways that are involved. Therefore, we set out to determine the *in vitro* requirements of the T cell “stop” signal linked to CXCR4.

## Results

### Monomeric vs Dimeric vs Polymeric CXCL12: T Cell Motility and Adhesion

To investigate the role of the polymeric CXCL12-coating of PDA cells in modulating T cell motility *in vitro*, we compared the effects of stimulating T cells with increasingly multimeric forms of the chemokine: monomer, dimeric (CXCL12)_2_-Fc fusion protein, and the tetramer comprised of the protein A complex with two (CXCL12)_2_-Fc fusion proteins (13). Monomeric CXCL12, which signals through CXCR4 *via* G protein-coupled receptor pathway, did not significantly affect the motility of the human T cells.

In contrast, an equivalent concentration of the dimeric (CXCL12)_2_-Fc fusion protein significantly reduced T cell motility, and a more highly polymeric form of CXCL12, which is (CXCL12)_2_-Fc complexed to protein A, resulted in a more pronounced cessation of T cell movement (Fig. 1a).

**Figure 1.**
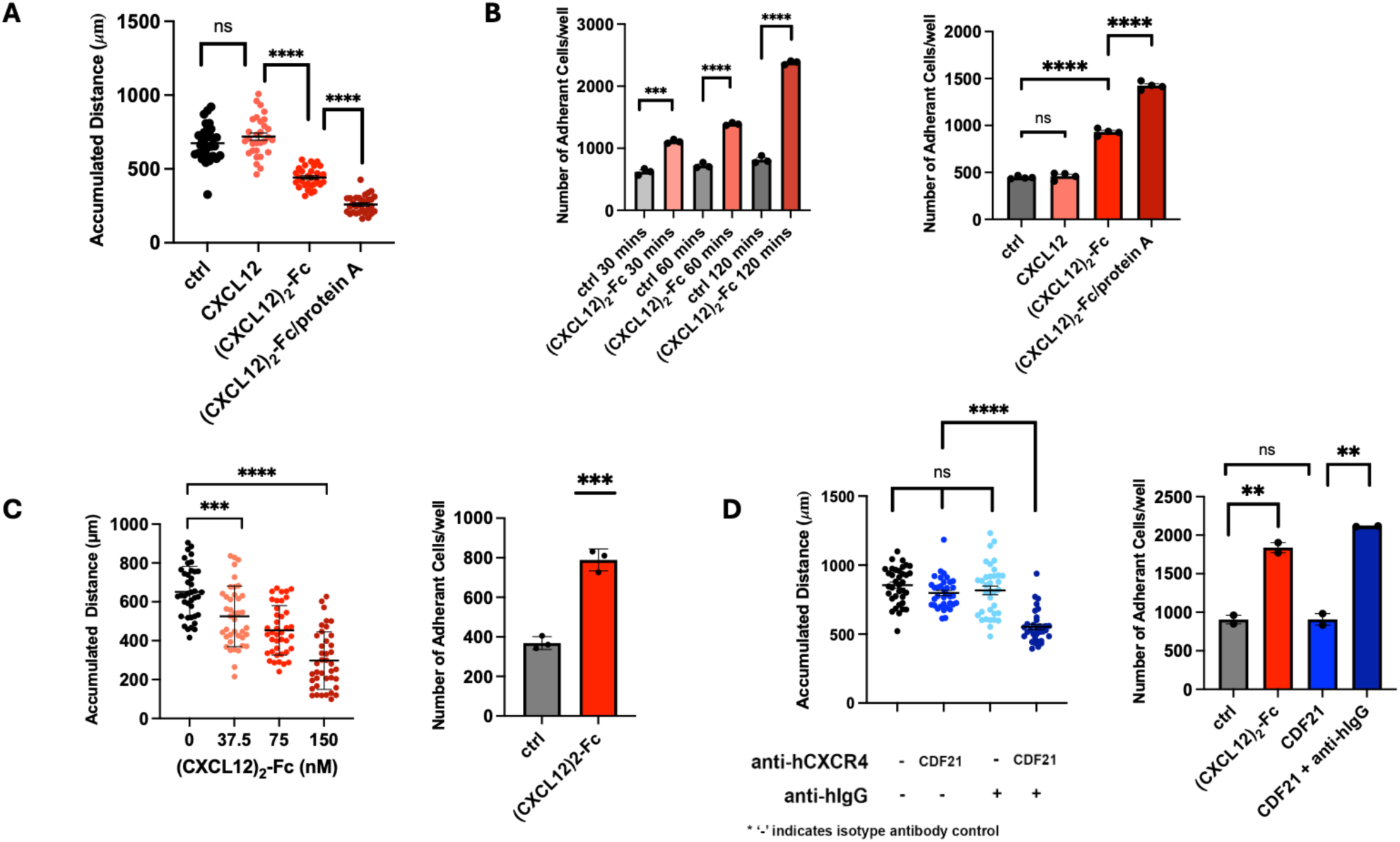
The e%ect of crosslinking CXCR4 on T cell motility and adhesion to fibronectin. **A.** HSB2 T cells were treated with 50 nM CXCL12 monomer, the (CXCL12)_2_-Fc fusion protein, and the (CXCL12)_2_-Fc/protein A complex, respectively, and motility of single T cells on a fibronectin-coated surface was measured for 2 hours (n = 30). **B.** HSB2 T cells were treated with 150 nM of the (CXCL12)_2_-Fc fusion protein for 1 hour then added to fibronectin-coated plates for di%erent time intervals. The plates were lightly washed after which the number adherent T cells were counted (n = 3). The treatments of the HSB2 T cells in panel A were applied to the adhesion assay (n = 4). **C.** Shown are the results of stimulating primary human e%ector T cells with incremental concentrations of the (CXCL12)_2_-Fc fusion protein on spontaneous motility (n = 40) and adhesion to fibronectin (n = 3). **D.** HSB2 T cells were incubated with CDF21, a human anti-human CXCR4 antibody, washed, and incubated with an isotype control rabbit IgG or a rabbit anti-human IgG antibody. The motility (n = 30) and adhesion (n = 2) of the HSB2 T cells were measured. Mean ± SEM; ns, not significant, **P < 0.01, ***P < 0.001, ****P < 0.0001, Student’s t test.

To examine whether the observed inhibition of T cell motility was caused by adhesion to fibronectin, we treated HSB2 T cells with the dimeric (CXCL12)_2_-Fc fusion protein, and measured the number of cells that remained on a fibronectin-coated surface after light washing. We observed significantly adhesion of the the T cells to the fibronectin-coated plates that increased in a time-dependent manner (Fig. 1b). We extended these observations by treating the HSB2 T cells with equivalent concentrations of the monomeric, dimeric, and polymeric forms of CXCL12, respectively, and measuring adherence to fibronectin. The results revealed that the more that CXCL12 was multimeric, the more the chemokine was able to enhance HSB2 T cell adhesion to fibronectin (Fig 1b).

It was important to demonstrate the results obtained with the HSB2 lymphoblastoid T cell line were also observed with primary effector human T cells, as would be present in the tumor miocroenvironment. For this purpose, we cultured normal peripheral blood human T cells with anti-CD3e and IL-2, and then rested the cells. When these T cells were stimulated with incremental concentrations of the (CXCL12)_2_-Fc fusion protein, they demonstrated a dose-related impairment of spontaneous motility and adhesion to the fibronectin-coated plates (Fig. 1c).

We hypothesized that the decreased T cell motility and increased adhesion occurring with multimeric CXCL12, and monomeric CXCL12, was caused by crosslinking of CXCR4. We tested this possibility by binding an anti-human CXCR4 human monoclonal antibody to HSB2 T cells, which does not stimulate the CXCR4 chemotactic response, and crosslinking the cell-bound antibody with a polyclonal anti-human IgG antibody. This mode of crosslinking CXCR4 caused similar decreases in HSB2 T cell spontaneous motility and increases in adhesion to fibronectin,as did polymeric (CXCL12)_2_-Fc/protein A complex (Fig. 1d).

### Phosphorylation of PTK2B and the CXCR4-Mediated “Stop” Signal

Recognizing that the crosslinking of CXCR4 and increased adhesion of T cells to fibronectin *in vitro* could represent an *in vivo* “stop” signal for the excluded T cells that were interacting with the polymeric CXCL12-coat in pancreactic cancer cells, we sought the basis for this response. Cellular adhesion to fibronectin is frequently mediated by the focal adhesion kinases, PTK2 and PTK2B (14). Since immunoblot analysis indicated that HSB2 T cells express low levels of PTK2 and higher levels of PTK2B (Fig. S1), we focused on whether crosslinked CXCR4 induced tyrosine phosphorylation of PTK2B. Upon treatment of the HSB2 T cells with (CXCL12)_2_-Fc fusion protein, we observed tyrosine phosphorylation of PTK2B within five minutes which persisted for at least one hour (Fig. 2a). Notably, using antibodies specific for the different phosphotyrosines of PTK2B, we were able to observe that this phosphorylation event occurred on tyrosine-579/580 of PTK2B rather than tyrosine-402 (Fig. 2b). We assessed the effects of monomeric, dimeric, and tetrameric CXCL12 stimulation of HSB2 T cells on tyrosine phosphorylation of PTK2B and demonstrated that only the dimeric and tetrameric forms induced phosphorylation of PTK2B (Fig. 2c), with tetrameric form of (CXCL12)_2_-Fc fusion/protein A complex inducing more intense tyrosine phosphorylation. We examined the functional requirement for PTK2B in the CXCR4-dependent motility and adhesion effects by treating HSB2 T cells with the kinase inhibitor, PF-4618433, that is selective for PTK2B (15). This treatment blocked the effects of the (CXCL12)_2_-Fc fusion protein on T cell motility and adhesion. Thus, PTK2B activation is an essential mediator of the “stop” signal induced by crosslinking CXCR4 (Fig. 2d).

**Figure 2.**
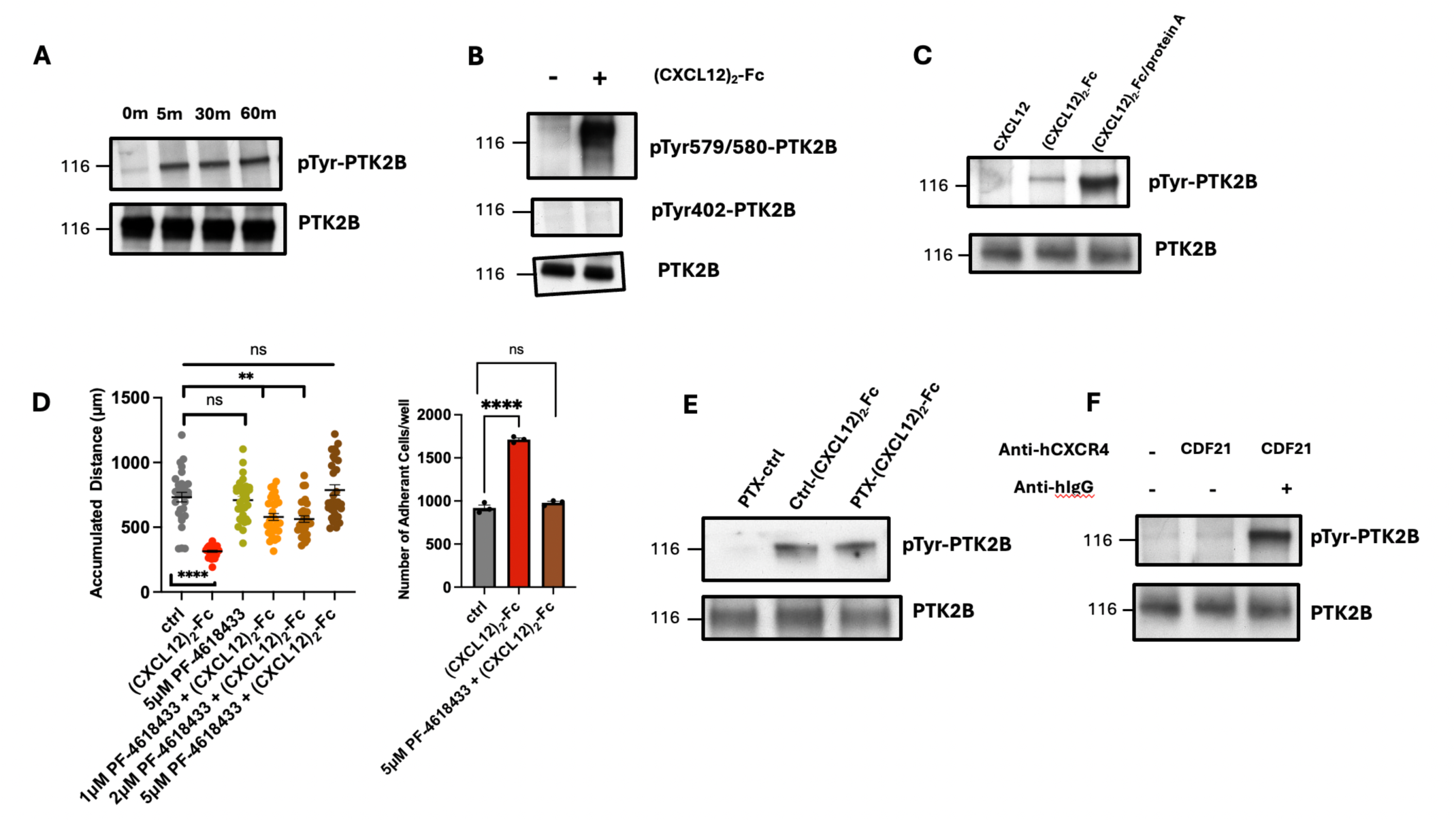
Tyrosine phosphorylation of PTK2B and crosslinking of CXCR4. **A.** HSB2 T cells were incubated with the (CXCL12)_2_-Fc fusion protein for timed intervals, after which PTK2B was immunoprecipitated from cell lysates, subjected to SDS-PAGE, and the gel was immunoblotted with antibodies to PTK2B and phosphotyrosine, respectively. **B.** HSB2 T cells were incubated with the (CXCL12)_2_-Fc fusion protein and cell lysates were subjected to SDS-PAGE and immunoblotting with antibodies to pTyr579/580 of PTK2B and pTyr402, respectively. **C.** HSB2 T cells were incubated with the monomeric CXCL12, dimeric CXCL12 and tetrameric CXCL12, respectively, and the cell lysates were analyzed as in panel A. **D.** HSB2 T cells were pre-incubated with the PTK2B inhibitor, PF-4618433, followed by incubation with the (CXCL12)_2_-Fc fusion protein, after which cell motility (n = 30) and adhesion to fibronectin (n = 3) were assessed. **E.** HSB2 T cells were subjected to treatment with PTX or bu%er control, followed stimulation with the (CXCL12)_2_-Fc fusion protein. The cell lysates were analyzed for tyrosine phosphorylation of PTK2B. **F.** HSB2 T cells were treated with CDF21 human anti-CXCR4 antibody followed by anti-human IgG antibody, and cell lysates were analyzed for tyrosine phosphorylation of PTK2B. Mean ± SEM; ns, not significant, **P < 0.01, ****P < 0.0001, Student’s t test.

Using the biochemical response of PTK2B to this manner of ligating CXCR4, we assessed the effects of pertussis toxin (PTX), which ADP-ribosylates Gα*i*, thereby inhibiting this pathway of CXCR4 signaling. The presence of PTX had no effect on PTK2B tyrosine phosphorylation, demonstrating that this step of the T cell "stop" signal that is elicited by crosslinking CXCR4 is distinct from the chemotactic response of CXCR4 to CXCL12 (Fig. 2e). This result is consistent with the capacity of the anti-CXCR4 monoclonal antibody, which does not elicit a chemotactic signal, to induce the tyrosine phosphorylation of PTK2B (Fig. 2f). Thus, crosslinking CXCR4 engages a distinct, non-chemotactic pathway when mediating arrested motility and enhanced adhesion to fibronectin.

### Integrins and TNFa in the Adhesion of T Cells to Fibronectin

Building on our observations of increased T cell adhesion to fibronectin, we investigated the possibility of an integrin-mediated response in this process. Flow cytometry analysis revealed that both HSB2 T cells and primary T cells express high levels of integrin subunits α4 and β1 (16). Inhibiting α4 integrin with a blocking antibody led to significant inhibition of the effect of the (CXCL12)_2_-Fc fusion protein on HSB2 T cell motility (Fig. 3a) and adhesion to fibronectin (Fig. 3b). In an attempt to sequence the cellular responses to crosslinking CXCR4, we pretreated HSB2 T cells with PTX and observed inhibition of adhesion to fibronectin induced by (CXCL12)_2_-Fc (Fig. 3c). As PTX had no effect on tyrosine phosphorylation of PTK2B, this response indicates that the activation of PTK2B by crosslinking CXCR4 is upstream of the activation of integrin α4.

**Figure 3.**
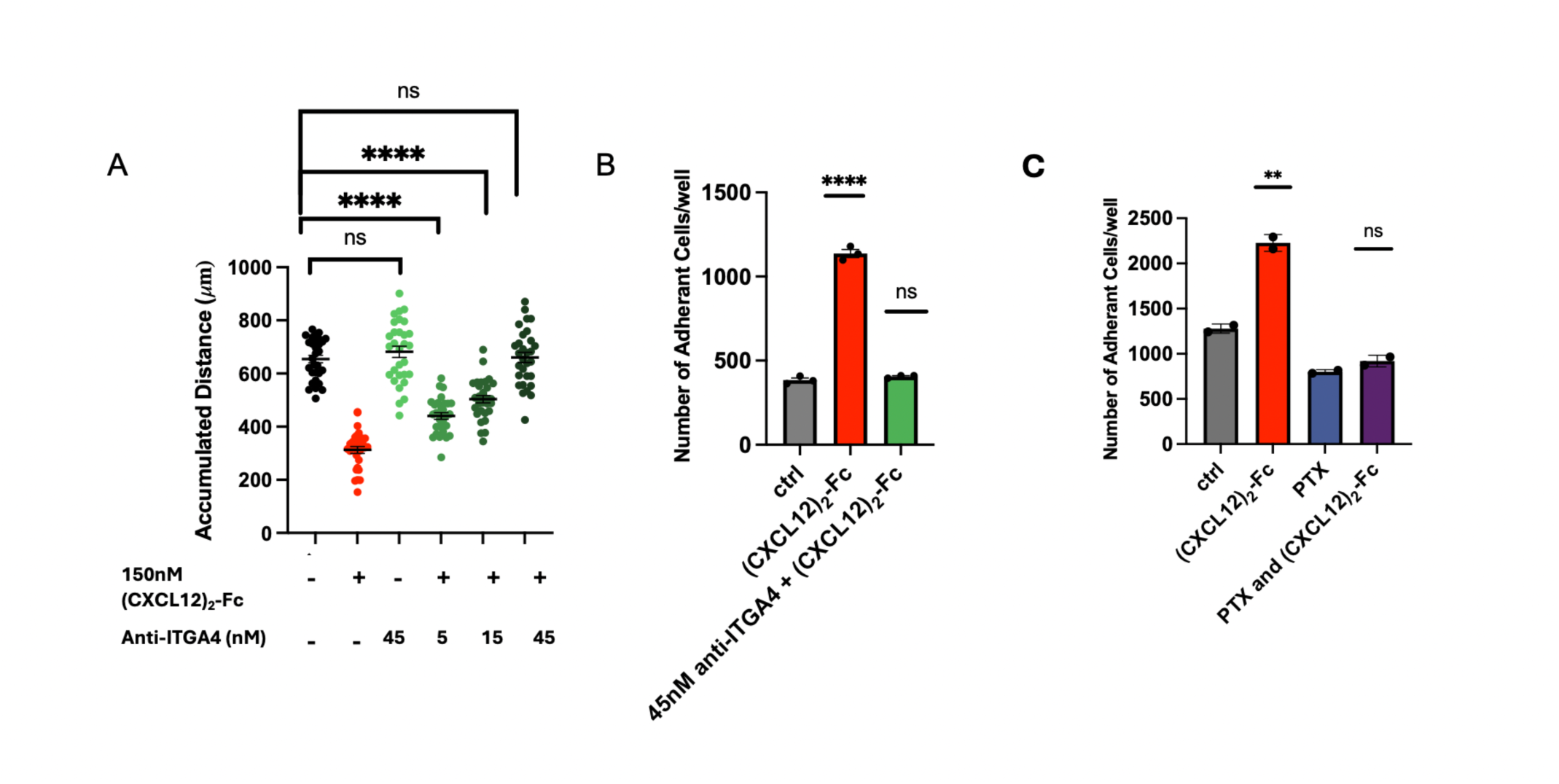
The role of integrins in T cell immotility and adhesion. **A.** HSB2 T cells were pretreated with the anti-ITGA4 antibody followed by treatment with the (CXCL12)_2_-Fc fusion protein, after which (**A**) cell motility (n = 30) and (**B**) adhesion to fibronectin (n = 3) were measured, respectively. **C.** HSB2 T cells were pretreated with PTX followed by treatment with the (CXCL12)_2_-Fc fusion protein, after which adhesion to fibronectin was measured in duplicates. Mean ± SEM; ns, not significant, **P < 0.01, ****P < 0.0001, Student’s t test.

TNFα and its receptors, TNFRSF1 and TNFRSF1B, may have a role in integrin-mediated T cell adhesion (17) and even a “stop” signal for T cells (18). HSB2 T cells spontaneously secrete TNFα (Fig. S2), and if this TNFα is neutralized by an anti-TNFα antibody, the effects of the (CXCL12)_2_-Fc fusion protein on T cell motility and adhesion are abolished (Fig. 4a). HSB2 T cells express mainly TNFRSF1B rather than TNFRSF1, suggesting that stimulation of TNFRSF1b promotes integrin-mediated adhesion to fibronectin (Fig. S3). We confirmed this conclusion by conducting assays with Jurkat T cells. These cells do not secrete TNFα or express TNFRSF1B, but does express TNFRSF1 (19). When provided with exogenous TNFα and stimulated with the (CXCL12)_2_-Fc fusion protein, the Jurkat T cells did not exhibit inhibited motility (Fig. S4). However, when the assay was performed with Jurkat T cells that expressed TNFRSF1B secondary to lentiviral transduction (Fig. 4b) and were treated with exogenous TNFα, they exhibited both motility inhibition (Fig. 4c) and increased adhesion to fibronectin (Fig. 4d) in response to the (CXCL12)_2_-Fc fusion protein. Therefore, TNFRSF1B signaling is necessary for the adhesion and motility effects triggered by crosslinked CXCR4.

**Figure 4.**
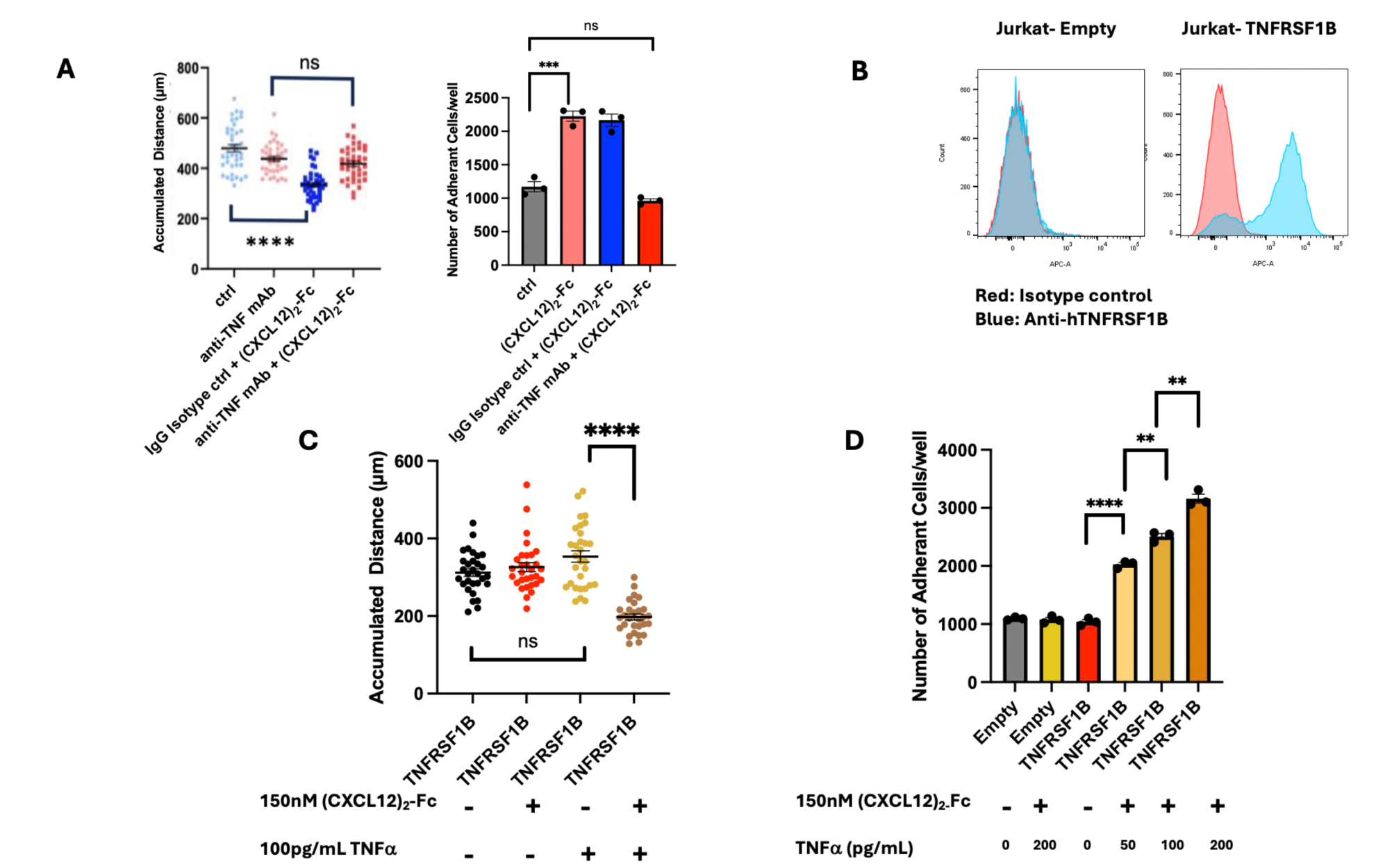
TNFα and TNFRSF1B signaling and T cell motility and adhesion. **A.** HSB2 T cells were pretreated with a neutralizing antibody to TNF followed by stimulation with the (CXCL12)_2_-Fc fusion protein. T cell motility (n = 40) and adhesion to fibronectin (n = 3) were then measured. **B.** Jurkat T cells were lentiviral vector expressing TNFRSF1B and empty vector control, respectively. Combinations of the (CXCL12)_2_-Fc fusion protein and TNFα were added to the Jurkat T cells and they were assessed for motility (n = 30) (**C**). **D.** Jurkat T cells that had been transduced with a lentiviral vector expressing *TNFRSF1B* or an empty vector control were incubated with increasing concentrations of TNFα, and adhesion of the T cells to fibronectin was measured (n = 3). Mean ± SEM; ns, not significant, *P < 0.05, **P < 0.01, ***P < 0.001, ****P < 0.0001, Student’s t test.

## Discussion

In this study, we identify a previously unrecognized mechanism that possibly underpins T cell exclusion in PDA. Our findings reveal that the key barrier to T cell migration may be an unique form of signaling that induced by crosslinking CXCR4 by physical coating of the cancer cells with complexes of KRT19 and CXCL12. *In vitro* this manifests as decreased spontaneous motility of T cells caused by their increased adhesion to fibronectin. While immune exclusion in PDA has been linked to multiple factors, including the protective stromal environment and immune checkpoint evasion, these findings suggest that crosslinking CXCR4 on T cells by the polymeric CXCL12 on cancer cells suppresses motility, leading to the exclusion of T cells.

We highlight the discovery of the CXCR4-mediated “stop” signal, a newly identified pathway that is distinct from typical chemotactic signals mediated by Gαi.

Unlike conventional CXCR4 signaling, which is dependent on G protein activation, this pathway relies on crosslinking CXCR4 in a G protein-independent manner. Our findings provide a molecular explanation for the long-recognized phenomenon of T cell exclusion by carcinoma cells, a key mechanism in immune evasion in PDA. The polymeric form of CXCL12, which is characteristic of the tumor’s protective coat, plays a critical role in this process. By inducing a potent G protein-independent stop signal, polymeric CXCL12 effectively halts T cell migration, leading to the exclusion of T cells from the tumor microenvironment. These insights reveal a novel mechanism by which the tumor actively prevents immune infiltration, highlighting the role of polymeric CXCL12 in immune evasion in PDA.

The adhesion of T cells to fibronectin plays a critical role in this process. Our findings show that exposure to dimeric and polymeric CXCL12 significantly increases T cell adhesion to fibronectin-coated surfaces, a key event in the inhibition of T cell motility. This adhesion response is distinct from the effects observed with monomeric CXCL12, suggesting that the polymeric form is far more potent in triggering adhesion and migration arrest.

Following these observations, we turned our attention to the signaling pathways that mediate the CXCL12-induced stop signal. Notably, dimeric and polymeric CXCL12 trigger rapid phosphorylation of PTK2B, a kinase well-known for its role in cell motility and adhesion. This phosphorylation event, which occurs in response to CXCR4 crosslinking by polymeric CXCL12, leads to T cell adhesion to fibronectin and directly contributes to the cessation of their movement. Our use of a specific PTK2B inhibitor rescued T cell motility, solidifying PTK2B phosphorylation as a central mediator of the stop signal and subsequent adhesion.

Furthermore, we demonstrate that TNFα and TNFRSF1B signaling are integral to the adhesion process. Soluble TNFα, secreted by both T cells and macrophages in the PDA microenvironment, and its binding to TNFRSF1B are required for the adhesion of T cells to fibronectin following PTK2B activation. This TNFα-TNFRSF1B binding leads to an integrin-dependent interaction between the T cell and fibronectin, promoting further adhesion and reinforcing the block to motility. Interestingly, while PTK2B phosphorylation occurs through a G protein-independent mechanism, the integrin involvement and subsequent adhesion to fibronectin are G protein-dependent, highlighting a bifurcated signaling pathway that governs T cell behavior in response to the tumor microenvironment. This division of signaling-G protein-independent PTK2B activation followed by G protein-dependent integrin activation-suggests that targeting both the CXCR4-PTK2B axis and the TNFα-TNFRSF1B-integrin pathway could be essential for reversing T cell exclusion in PDA.

In conclusion, our findings provide a comprehensive understanding of the molecular mechanisms that facilitate immune evasion in PDA. The interplay between CXCL12-CXCR4 signaling, PTK2B phosphorylation, and TNFα-TNFRSF1B signaling constitutes a multifaceted barrier to T cell migration and infiltration. These insights have significant therapeutic implications, suggesting that disrupting the CXCL12-CXCR4-PTK2B and TNFα-TNFRSF1B signaling pathways may restore T cell motility and improve immune responses against PDA. Given the limited success of current immune therapies, particularly immune checkpoint inhibitors, targeting these pathways could represent a promising strategy to enhance immune infiltration and therapeutic outcomes.

## Methods

### Fibronectin ibidi Assay for Cell Migration

Cell migration was assessed using an Ibidi 6-well chamber slide (Ibidi, 80626). Cells were treated with 150 nM (CXCL12)_2_-Fc (Sinobiological, 10118-H01H) or vehicle control for 1 hour at 37°C in chemotaxis buffer (RPMI-1640 supplemented with 0.5% BSA and 20 mM HEPES). For experiments involving inhibitors or antibodies*, cells were pretreated for 15 minutes at 37°C before the addition of (CXCL12)_2_-Fc. Fibronectin (Sigma) was diluted to 75 µg/mL in PBS containing 2 mM CaCl₂ and 2 mM MgCl₂ and added to each channel of the Ibidi chamber slide (40 µL per channel). The plate was incubated at room temperature for 1 hour to allow fibronectin coating. Excess fibronectin was aspirated, and the channels were washed once with 40 µL of TBS, followed by another wash with 40 µL of chemotaxis buffer. Cells were then diluted 1:3 in chemotaxis buffer to a final concentration of 3.33 × 10⁵ cells/mL. A total of 40 µL of the cell suspension was added to each channel, avoiding the formation of air bubbles. The chambers were sealed with scotch tape to prevent evaporation and maintain an airtight system. The Ibidi plate was placed into a chamber maintained at 37°C with 5% CO₂ and allowed to acclimate for 15 minutes before imaging. Cell migration was monitored for 2 hours, with images taken every 3 minutes using a ZEISS Observer microscope. Cell movements were manually tracked using ImageJ software and analyzed with the Ibidi Chemotaxis and Migration Tool (https://ibidi.com/chemotaxis-analysis/171-chemotaxis-and-migration-tool.html).

*.TNF neutralizing ab (Cell Signaling, 7321), PF-461833 (MedChemExpress, HY-18312), Integrin Alpha 4 Inhibitor (Biotechne, MAB10603).

### Cell Adhesion Assay on Fibronectin-Coated 96-Well Plates

Fibronectin-coated 96-well plates were prepared by incubating wells with human plasma fibronectin (50 µg/mL) in PBS supplemented with 2 mM CaCl₂ and 2 mM MgCl₂ overnight at 4°C. Non-specific binding sites were blocked by incubating control wells with 5% BSA in PBS++ at 4°C overnight. The next day, excess fibronectin was aspirated, and the wells were washed once with TBS. Blocking was completed by incubating the wells with 100 µL of 2% non-fat milk in TBS at room temperature for 30 minutes. After blocking, the wells were washed three times with TBS, and the treated cells, resuspended in appropriate culture medium, were added to each well at 100 µL per well. The plate was incubated at 37°C for 1 hour to allow cell adhesion. Following the incubation, the plate was inverted onto a paper towel and allowed to sit at room temperature for 20 minutes to facilitate the removal of non-adherent cells. The wells were washed three times with TBS to remove any residual non-adherent cells. Adherent cells were then imaged using a light microscope for assessment of cell adhesion and morphology.

### Polymeric (CXCL12)_2_-Fc Preparation

Protein A (Pierce, 21184) was diluted to a final concentration of 200 nM and combined with an equivalent volume of 400 nM (CXCL12)_2_-Fc to achieve a final concentration of 100 nM protein A and 200 nM (CXCL12)_2_-Fc. A 1:2 ratio, determined to be the saturation point for (CXCL12)_2_-Fc binding to protein A, was used to ensure complete saturation, resulting in a final polymer concentration of 100 nM. The mixture was incubated at 37°C for 1 hour.

### Immunoprecipitation of Phosphorylated PTK2B

Human CCRF-HSB-2 T lymphoblast cells (ATCC, CCL-120.1) were maintained in RPMI-1640 medium (Corning) supplemented with 10% FBS (Seradigm), 100 U/mL penicillin, and 100 µg/mL streptomycin, at a density of 1 × 10⁶ cells/mL. On the day of the experiment, cells were washed with PBS, counted, and resuspended in chemotaxis buffer (RPMI-1640 supplemented with 20 mM HEPES, pH 7.5, and 0.5% BSA) at a density of 2.67 × 10⁶ cells/mL. A total of 4 × 10⁶ cells were plated per treatment in 1.5 mL of chemotaxis buffer. After treatment, cells were collected by centrifugation at 300 x g for 5 minutes, and the supernatant was discarded. The resulting pellet was resuspended in 75 µL of lysis buffer (Pierce RIPA (Thermo, 89900) + Pierce Proteinase and Phosphatase inhibitor cocktail (Thermo, 78447)) and incubated on a rotating platform at 4°C for 30 minutes. The lysate was then centrifuged at 16,000 x g for 10 minutes at 4°C. The supernatant was transferred to a new tube, and 200 µg of total lysate was combined with 20 µg of primary antibody (Cell Signaling, 3285). The volume was adjusted to 200 µL with lysis buffer, and the mixture was rotated overnight at 4°C. Dynabead Protein G beads (Thermo, 10003D) were incubated overnight at 4°C in lysis buffer supplemented with 1% BSA. The lysate and beads were then combined and incubated on rotation at 4°C for 2 hours. Afterward, the beads were washed four times with lysis buffer, resuspended in sample buffer, and boiled at 100°C for 10 minutes. The lysate was loaded into SDS-PAGE wells and analyzed by Western blot using anti-phosphotyrosine antibody, clone 4G10 (Sigma, 05-321X).

### Human Primary T Cell Isolation and Activation

A human blood sample was collected from a consenting donor, following ethical guidelines. CD3+ T cells were isolated using the Direct Human T-Cells Isolation Kit (Stemcell Technologies). The isolated naïve T cells were then activated by incubation with Dynabead Human CD3/CD28 Activating Beads (Thermo, 11131D), along with human interleukin-2 (R&D, BT-002-010) in complete T-cell medium for a minimum of 2 days. After activation, the activation beads were removed, and the activated T cells were resuspended in complete T-cell medium containing human interleukin-2. The cells were subsequently cultured for an additional 3 days.

### Transduction of Jurkat Cells with TNFRSF1B

LentiCas9_BSD was used as the backbone vector, and TNFRSF1B was inserted using AgeI and BamHI restriction sites. In-fusion cloning was employed to generate the construct EF1a-AgeI-TNFRSF1B-BamHI-P2A-BSD. The vector was transfected into the 293FT cell line using the lenti X single-shot transfection method, and the virus was collected 48 hours post-transfection. The collected virus was concentrated 10x using the lenti X concentrator. Jurkat cells (ATCC, TIB-152) were prepared at approximately 1 million cells per well in a 6-well plate and treated with 20 µg/mL polybrene. Virus titration was performed using serial dilutions (500 µL, 250 µL, 125 µL, 62.5 µL, 31.25 µL, and 0 µL) for transduction of Jurkat cells. After overnight transduction, the cells were washed and resuspended in fresh RPMI medium. Seventy-two hours post-transduction, Jurkat cells were selected with 7.5 µg/mL blasticidin for stable cell line establishment.

### Flow Cytometry

Cells were diluted to a concentration of 2 x 10^6^ cells/mL in appropriate medium. After being treated, 100 µL of each sample was transferred into a round-bottom 96-well plate and centrifuged at 2,000 rpm for 2 minutes to pellet the cells. The supernatant was discarded, and the cells were resuspended in 200 µL of FACS buffer (Thermo, 00-4222-26) and stained with various antibodies* for 30 minutes at 4°C. The cells were centrifuged and washed with 200uL FACS buffer three times to remove residual components. The cells were then resuspended a final time in 200 µL of FACS buffer and pipetted through the cell strainer cap of a flow tube (Fisher). To each tube, 400 µL of FACS buffer was added, bringing the final volume to 600 µL. DAPI (Thermo, R37606) was used for live-cell gating. Data acquisition was performed using a BD LSRFortessa cell analyzer with Diva software 8.0, and subsequent analysis was conducted using FlowJo

*Anti Human TNF-alpha antibody (biolegend, 502913), Isotype control (biolegend, 400119); Anti Human TNFRSF1A (biolegend, 369905), Isotype control (biolgend, 400221); Anti Human TNFRSF1B (biolegend, 358403), Isotype control (biolegend, 400507)

### ELISA

HSB2 T cells were treated with 150 nM (CXCL12)₂-Fc or vehicle control for various time points at 37°C in chemotaxis buffer (RPMI-1640 supplemented with 0.5% BSA and 20 mM HEPES). Following incubation, cells were centrifuged at 300 × g for 5 minutes at room temperature. The supernatant was collected and TNFα concentration was determined using a human TNF alpha ELISA kit (Thermo, KHC3011). The assay was performed according to the manufacturer’s instructions, and TNFα concentrations were quantified using a plate reader.

## Data Availability Statement

All study data are included in the manuscript and/or SI Appendix

## Author contributions

J.P., J.Y., S.K., Z.K. and D.T.F. designed research; J.P., J.Y., S.K., and Z.K. performed research; J.P., J.Y., S.K., Z.K., and D.T.F. analyzed data; and J.P. and D.T.F. wrote the paper.

## Acknowledgements

We acknowledge the contributions of the CSHL facilities for flow cytometry and microscopy. This work was supported by The Lustgarten Foundation (D.T.F) and Northwell Health (D.T.F).

## Abbreviations

(PDA): Pancreatic Ductal Adenocarcinoma
PTK2B: Proline-rich tyrosine kinase 2

