## Supplemental Information for "Crosslinked CXCR4 Signals Decreased Motility and Increased Adhesion of T Cells"

### Supporting Information Appendix

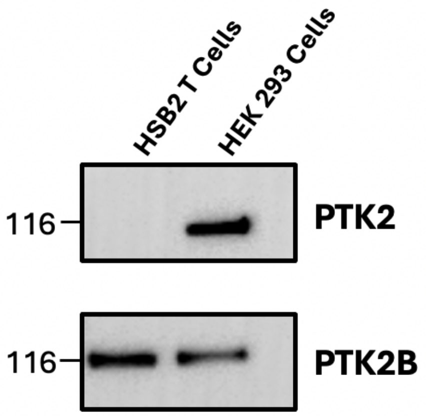


**Supplemental Figure 1.** HSB2 T cell lysates were subjected to SDS-PAGE followed by immunoblotting for PTK2 and PTK2B. HEK 293 cells were used as a positive control for PTK2 and PTK2B expression.

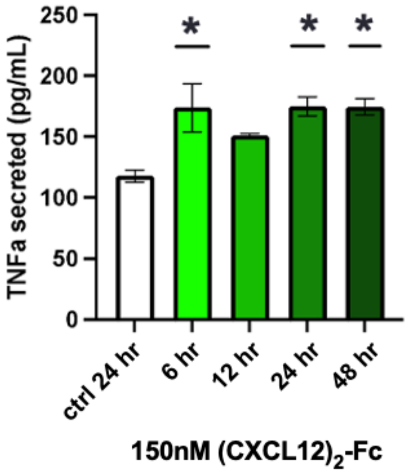


**Supplemental Figure 2.** HSB2 T cells were incubated with or without the $(CXCL12)_2$-Fc fusion protein for timed intervals. The supernatants were collected, and TNFα concentrations were measured by ELISA (n = 3). Mean ± SEM; *P < 0.05. Student's t test.

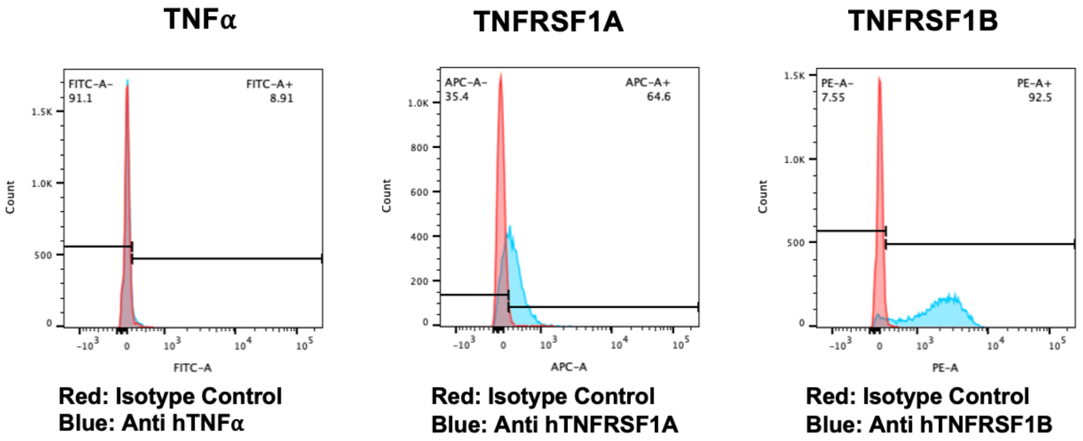


**Supplemental Figure 3.** The expression of membranous TNFα, TNFRSF1A, and TNFRSF1B on HSB2 T cells were measured by flow cytometry.

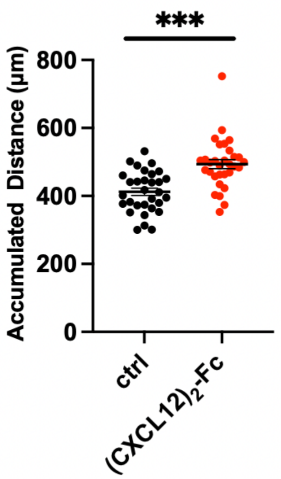


**Supplemental Figure 4.** Jurkat T cells were treated with TNFα, stimulated with the $(CXCL12)_2$-Fc fusion protein, and assessed for motility (n = 30). Mean ± SEM; ***P < 0.001, Student's t test.
